# The pace of life: Time, temperature, and a biological theory of relativity

**DOI:** 10.1101/609446

**Authors:** Anna B. Neuheimer

## Abstract

For living things, time proceeds relative to body temperature. In this contribution, I describe the biochemical underpinnings of this “biological time” and formalize the Biological Theory of Relativity (BTR). Paralleling Einstein’s Special Theory of Relativity, the BTR describes how time progresses across temporal frames of reference, contrasting temperature-scaled biological time with our more familiar (and constant) “calendar” time measures. By characterizing the relationship between these two time frames, the BTR allows us to position observed biological variability on a relevant time-scale. In so doing, we are better able to explain observed variation (both temperature-dependent and -independent), make predictions about the timing of biological phenomena, and even manipulate the biological world around us. The BTR presents a theoretical framework to direct future work regarding an entire landscape of fundamental biological questions across space, time and species.

## Introduction

The pace of life (e.g. time to stage, size, or lifespan) controls when biological players are observed, and how they interact with the environment and each other. Due to our “warm-blooded bias”, we experience this pace of life (biological time) as relatively constant (i.e. equivalent to measured “calendar” time), when in fact biological time accumulates relative to body temperature. Because of the biochemistry underlying biological processes such as growth, development, muscle contraction, even neural firing rates, biological time scales relative to the temperature experienced by the organism. Over a mid-range of temperatures, this means that biological time slows at cooler temperatures and speeds at warmer (but see Why biological time scales with temperature).

In this contribution, I formalize the relative nature of biological time as the Biological Theory of Relativity (BTR). Similar to Einstein’s Special Theory of Relativity (STR, see How biological time scales with temperature), this BTR allows us to relate time-scales across different frames of reference (i.e. biological vs. calendar). I present the foundations of the BTR, highlighting parallels to the STR. I show how using the BTR can help explain variation in biological observations over space and time, improve our predictions of how populations and ecosystems will behave in the future, and even allow us to manipulate the biological world around us. I finish by describing the research question landscape presented by the BTR, offering directions for future research that will allow us to exploit the BTR to move nimbly across temporal frames of reference to explain observations and make predictions about biological phenomena across space and time.

### Why biological time scales with temperature

At the heart of biological processes are chemical reactions that break down (e.g. digestion), build up (e.g. growth), or transform (e.g. movement), etc.

The rate of these reactions depend on available kinetic energy which signifies e.g. how likely and often substrates will meet. Biological reactions are often helped along by enzymes, proteins which increase the likelihood or speed of reactions, e.g. by increasing the probability of substrates meeting. That said, the rate of reactions overall depends on the kinetic energy of the system.

Kinetic energy is measured via the organism’s relevant body temperature. At lower temperatures, there is relatively little kinetic energy, and reaction rates proceed slowly (Fig. 1). At higher temperatures, there is more kinetic energy and reaction rates proceed more quickly. At even higher temperature, reaction rates decline. One reason for this decline is the denaturation of the enzymes involved in reaction rates. This pattern of increased reaction rates at increased temperatures, up to a maximum, is formalized in a relationship often termed the thermal performance curve, or TPC (Fig. 1).

**Fig. 1:**
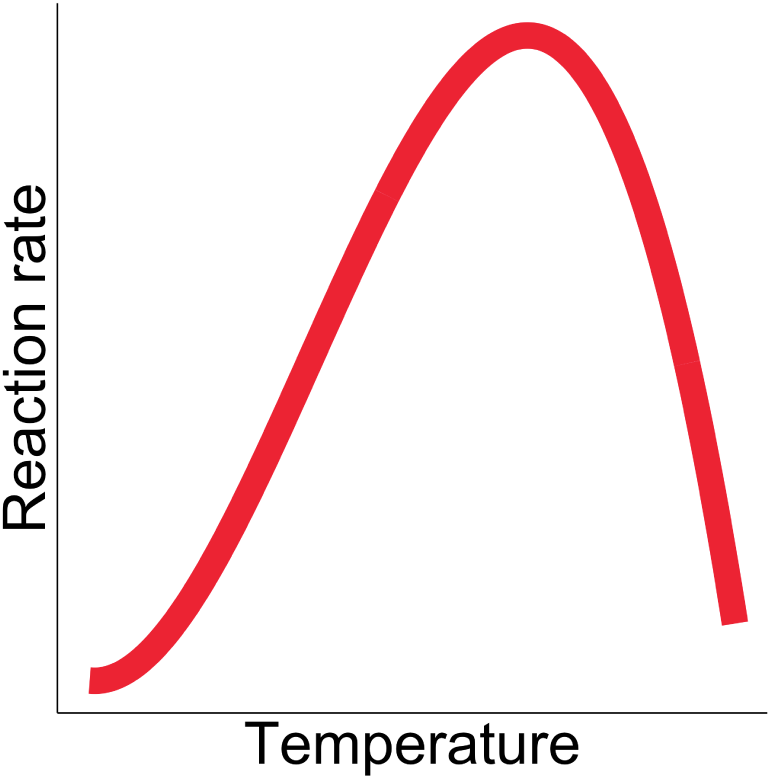
The thermal performance curve (TPC): The TPC describes how biological reaction rates respond to temperature with rates increasing with temperature up to a maximum.

While the TPC can be thought of for a particular physiological reaction rate, it is often measured at the level of individuals or populations where the dependent axis is e.g. growth or development rate. This latter approach implicitly includes additive and trade-off effects of all underlying reaction rates, e.g. compound transport, bond making or breaking, waste removal, etc. involved in the observed rate (e.g. growth rate). Characterizing the TPC and how it varies across processes, individuals, populations, etc. is fundamental as it determines how time scales with temperature - as I will discuss in the next section (see also Mapping out the BTR research landscape).

### How biological time scales with temperature

Biological time progresses relative to body temperature with biological time scaling relative to calendar time via the TPC. The BTR describes how this effect changes the temporal frame of reference experienced by biology in a manner similar to Einstein’s STR which allows us to compare measures of time across frames of reference moving at different speeds (Fig. 2).

**Fig. 2:**
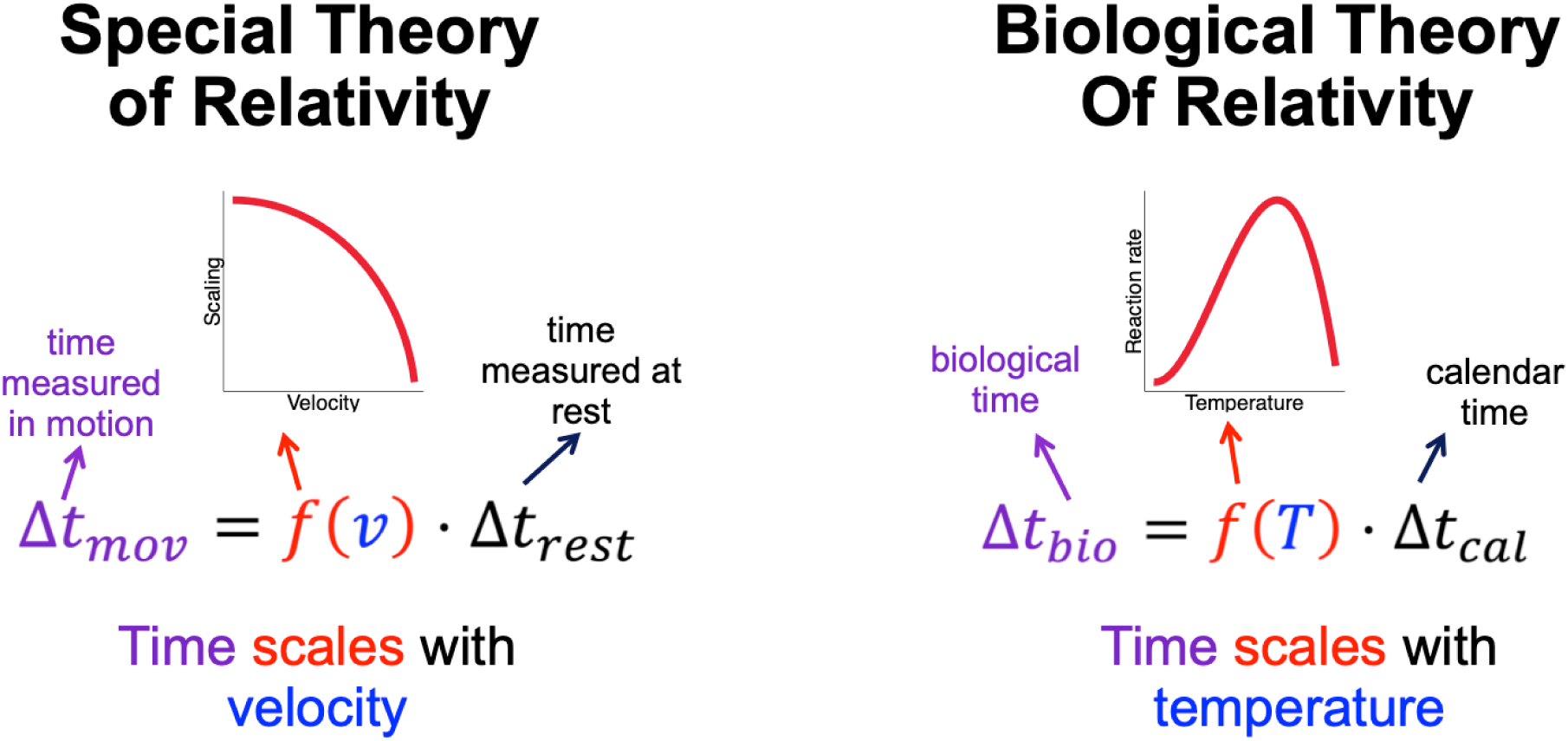
Parallels between the Special Theory of Relativity and the Biological Theory of Relativity. See text for description.

The STR describes how time scales with velocity:

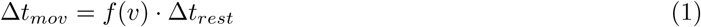

where Δ*t*_*mov*_ is the ellapsed time in the moving frame determined by the ellapsed time at rest (Δ*t*_*rest*_) and a velocity (*v*) scaling (*f* (*v*)), i.e. the Lorentz factor 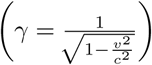 where *c* is the speed of light in a vacuum.

Similarly, the BTR describes how biological time scales with temperature as:

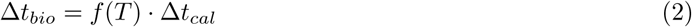

where Δ*t*_*bio*_ is biologically relevant elapsed time (i.e. in the biological frame of reference) determined by the elapsed “calendar” time (Δ*t*_*cal*_) scaled with the relevant body temperature (*T*) via the TPC (*f* (*T*)). Both body temperature (*T*) and the TPC (*f* (*T*)) may change in time & space (including across individuals). I will cover what happens when the TPC scaling changes in Mapping out the BTR research landscape below, but here let’s expand eqn (2) to allow for changes in body temperature over time (*T* (*t*)):

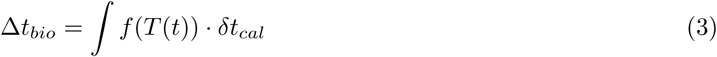

We can use eqn (3) to explore how biological age emerges relative to calendar age for warm-vs. cold-blooded organisms. **Warm-blooded** organisms (also called endotherms) are those animals that keep their temperature relatively constant regardless of external environmental temperatures (Fig. 3a). Here I consider those with the active ability to internally regulate their temperature but also discuss below behavioural temperature modifications and regional endotherms (see further in section). Because body temperatures remain constant, reaction rates are constant and, thus, the scaling of biological time (via the TPC) is constant for warm-blooded organisms (Fig. 3b).

**Fig. 3:**
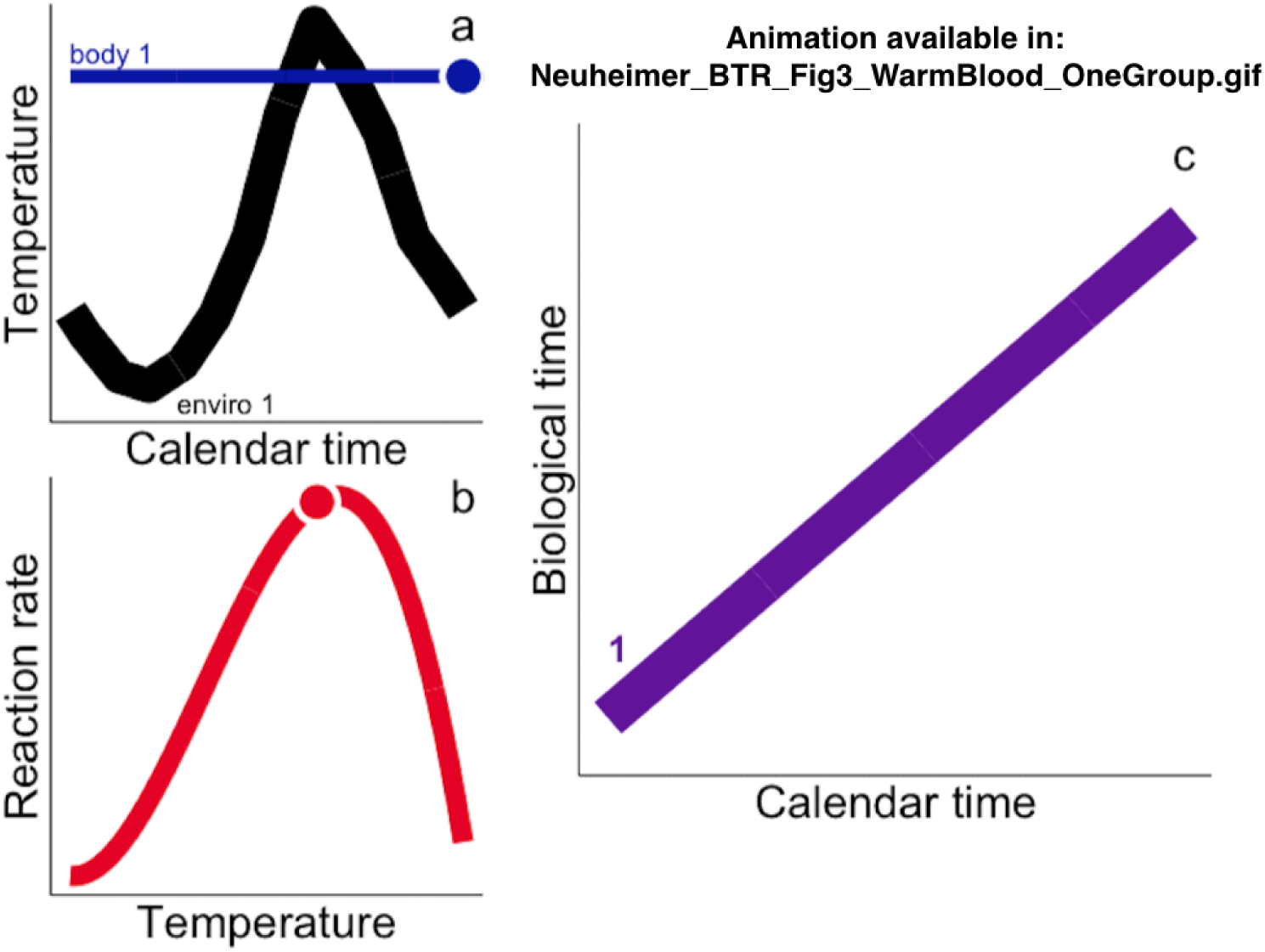
Illustrating biological time evolution for warm-blooded organisms. a) Warm-blooded organisms keep body temperatures (blue) constant relative to environmental temperatures (black). b) Constant body temperatures translates to constant reaction rates (time scaling) as sampled via the TPC. c) As a result, biological time accumulates at a constant rate relative to calendar time for warm-blooded organisms.

This trait allows warm-blooded organisms “control” over biological time, allowing biological time to accumulate at a constant, predictable rate (Fig. 3c) e.g. offering reliable performance for catching prey in cold weather [1], or allowing individuals to exploit large habitat ranges (e.g. humpback whales migration routes of up to 8400 km and 25°C change in habitat temperatures between feeding and birthing areas; [2], [3], [4]). Of course, the benefits of being warm-blooded come at a literal cost as basal metabolic rates in warm-blooded animals are 5 to 10 times higher than cold-blooded organisms of a similar size [5], [6]. The cost of actively maintaining your body temperature will depend on an organism’s surface area and heat conduction of the surrounding medium. One needs only to compare the smallest size of warm-blooded animals (mammals) on land (etrsucan shrew, *Suncus etruscus*, ∼0.1g) vs. in the water (harbor porpoise calf, ∼ 10kg) to see how thermoregulation expense can limit the minimum viable size across habitats depending on the heat conduction of the medium (i.e. air vs. water; [7]).

Now let’s consider how biological time accumulates for **cold-blooded** organisms (or ectotherms). Here I will use an example of an organism living in a seasonal temperature environment, though these ideas scale to other environments as well (scaling all the way back to the warm-blooded example when environmental temperatures are constant). As the environmental temperature varies over the year so too will body temperature (Fig. 4a), and thus the rates of the underlying reactions, according to the TPC (Fig. 4b). As such, biological time progresses differently than calendar time (Fig. 4c): as changing body temperature means the biological time scaling (via TPC) varies over the year, biological time accumulates more or less slowly when compared to constant calendar time.

**Fig. 4:**
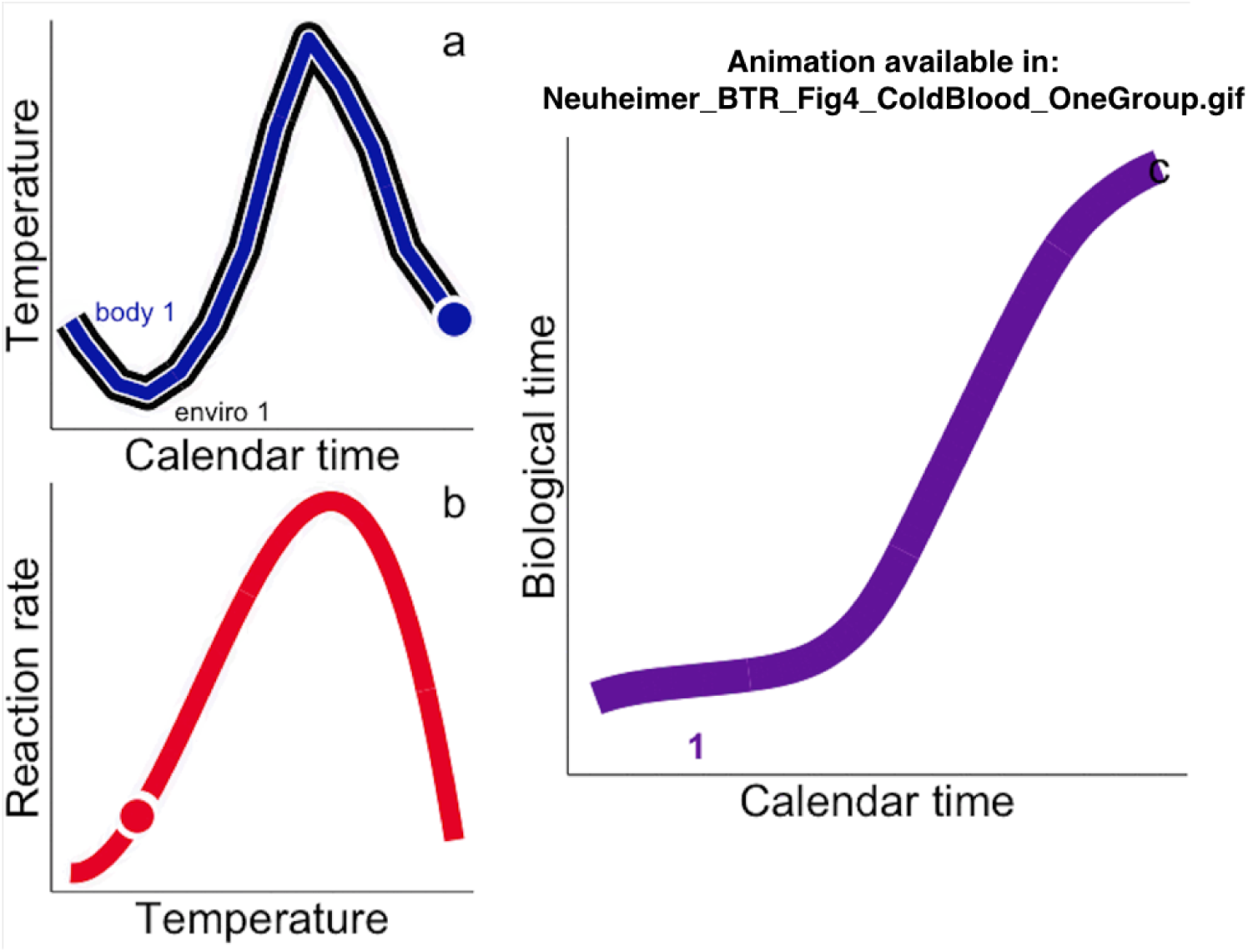
Illustrating biological time evolution for cold-blooded organisms. a) Body temperature (blue) of cold-blooded organisms track environmental temperatures (black). b) The TPC describes how these variable body temperatures result in variable reaction rates (time scaling). c) As a result, the accumulation of biological time varies with respect to calendar time for cold-blooded organisms.

Biological time for cold-blooded organisms accumulates not constantly then, but relative to environmental temperatures. To contrast with the warm-blooded animals above, this cold-blooded physiology is much cheaper to operate but means a cold-blooded organism’s clock is at the mercy of the environment. Cold-blooded organisms do have some tricks to get around this constraint. Organisms can change their body temperature by changing their spatial distribution (horizontally or vertically) and thus the environmental temperatures to which they are exposed (“behavioural thermoregulation”). Examples of this are many and include hibernating (diapausing) zooplankton (e.g. copepods) that overwinter in cool, deep waters minimizing environmental temperatures and effectively “slowing down” biological time during food-poor winter months [8], Atlantic stingrays that move to warmer waters when pregnant, speeding up the calendar time required for incubation [9], and whale sharks moving to cooler deeper waters to slow digestion time and likely increase nutrient absorption [10]. Alternatively, animals may adjust the ambient temperature of specific tissues to change the scaling of biological time (i.e. “regional endothermy”). An example of this is bluefin tuna (*Thunnus thynnus*) that heats swimming muscle tissue 10 to 20°C higher than the surrounding water [11] allowing for quicker rates of muscle contraction and more swimming power [12], [4].

### Using the biological theory of relativity

The usefullness of the BTR comes from an understanding of how biological time progresses relative to calendar time (e.g. Fig. 3c & 4c). Characterizing how biological time shapes what we see when we see it allows us to explain, predict, and even manipulate the world around us.

#### Explaining the world around us

First, let’s consider biological observations made in calendar time (e.g. size-at-age, time-to-stage) and how they might translate into biological time for warm-vs. cold-blooded organisms. With **warm-blooded** animals, body temperature remains constant from year to year (or location to location) despite possible changes in environmental temperature (Fig. 5a). This, combined with a TPC that is similar in space and time (but see Exploring the biology landscape on biological time) means the time scaling moving from calendar to biological time is constant (Fig. 5b) and biological time accumulates in a similar way across the groups (e.g. cohorts or locations, Fig. 5c).

**Fig. 5:**
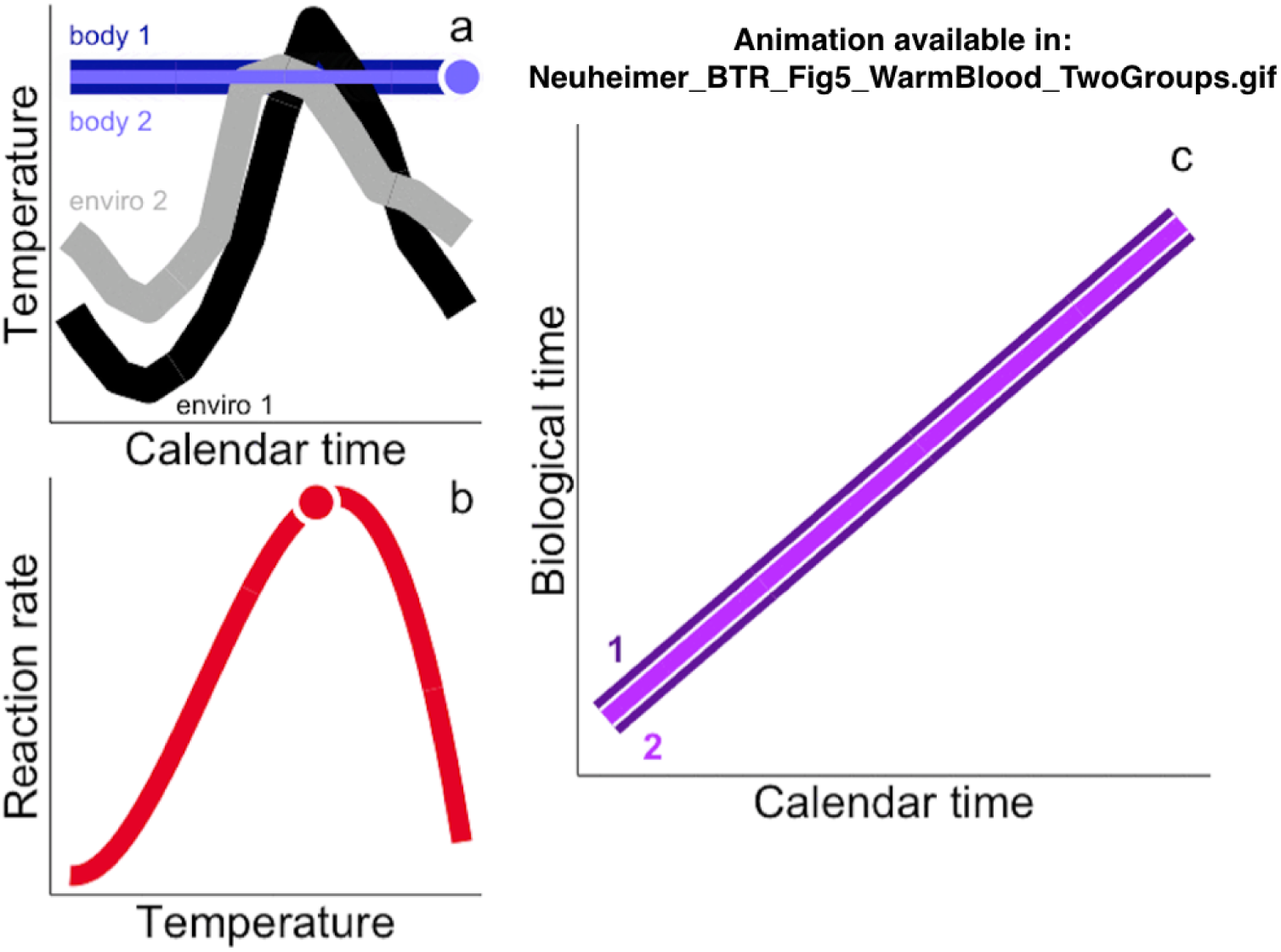
How biological time compares across groups for warm-blooded organisms. a) Environmental temperatures may change over space or years (grey vs. black) but warm-blooded organisms keep body temperatures constant (light vs. dark blue). b) Constant body temperatures translates to constant reaction rates and thus the time-scaling as sampled via the TPC stays the same across groups. c) As a result, biological time accumulates at a constant rate relative to calendar time for warm-blooded organisms and this evolution is similar across groups (light vs. dark purple).

This means that, as long as the TPC remains constant, biological observations measured in calendar time are arguably comparable across space and years for warm-blooded animals because the scaling to biological time is constant (Fig. 6). For example, we might compare the size at 5 months (calendar time) across populations or cohorts, comfortable in the idea that our calendar estimate of 5 months is comparable in both calendar and biological time-scales across the samples (Fig. 6). As warm-blooded animals then, we have a relatively easy time moving between the calendar and biological time-frames, e.g. comparing sizes of children of the same calendar age, or the calendar time to a developmental milestone such as a first tooth. These time-scales are comparable because we maintain a relatively constant body temperature. There are, of course, the potential that interesting, temperature-independent variability exists among warm-blooded animals (e.g. due to food consumption, genetics, etc.), but our use of calendar time as an approximate biological time-scale does not get in the way of identifying this interesting, temperature-independent variability.

**Fig. 6:**
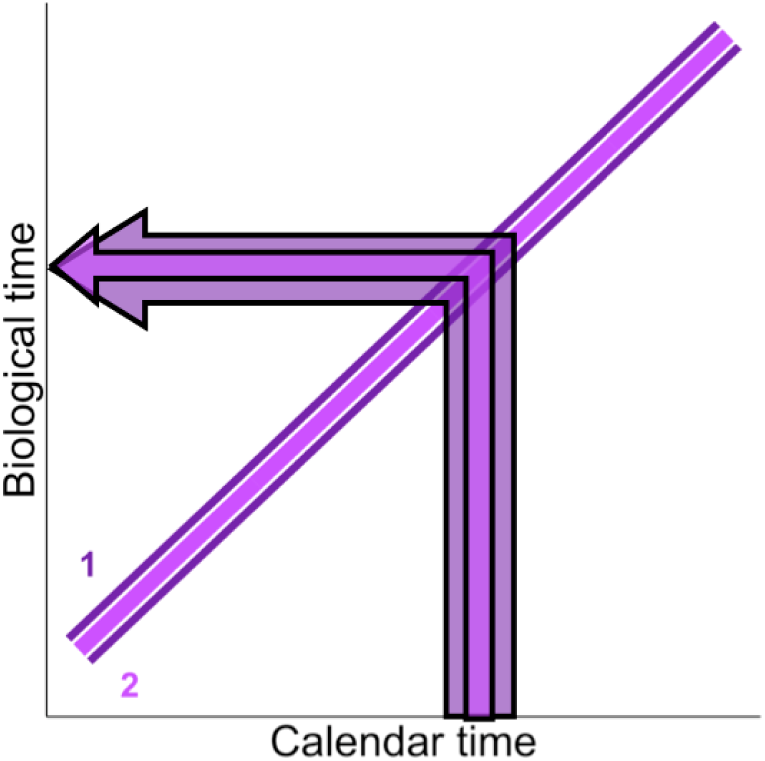
Calendar time observations can be directly compared across groups (e.g. cohorts or locations, light purple vs. dark purple) of warm-blooded organisms as the scaling between biological time and calendar time is held constant via constant body temperature.

Now, let’s contrast this with the accumulation of biological time by **cold-blooded** organisms living in different locations or years. Variations in the environmental temperatures across groups translate to variations in body temperature for cold-blooded organisms (Fig. 7a). These temperature variations along with the time-scaling as described by the TPC (Fig. 7b) result in biological time trajectories that emerge differently across groups in time or space (Fig. 7c). Thus, comparisons of observed timings for cold-blooded organisms on calendar time can be misleading when cold-blooded organisms occupy different thermal environments across space or time (Fig. 8). For example, comparing observations made at the same calendar time (e.g. size at 5 months) are difficult because the same point in calendar time (5 months) corresponds to a different amount of ellapsed biological time across the groups (Fig. 8, where an organism from group 2 would be ∼50% older in biological time than that from group 1 despite being the same calendar age).

**Fig. 7:**
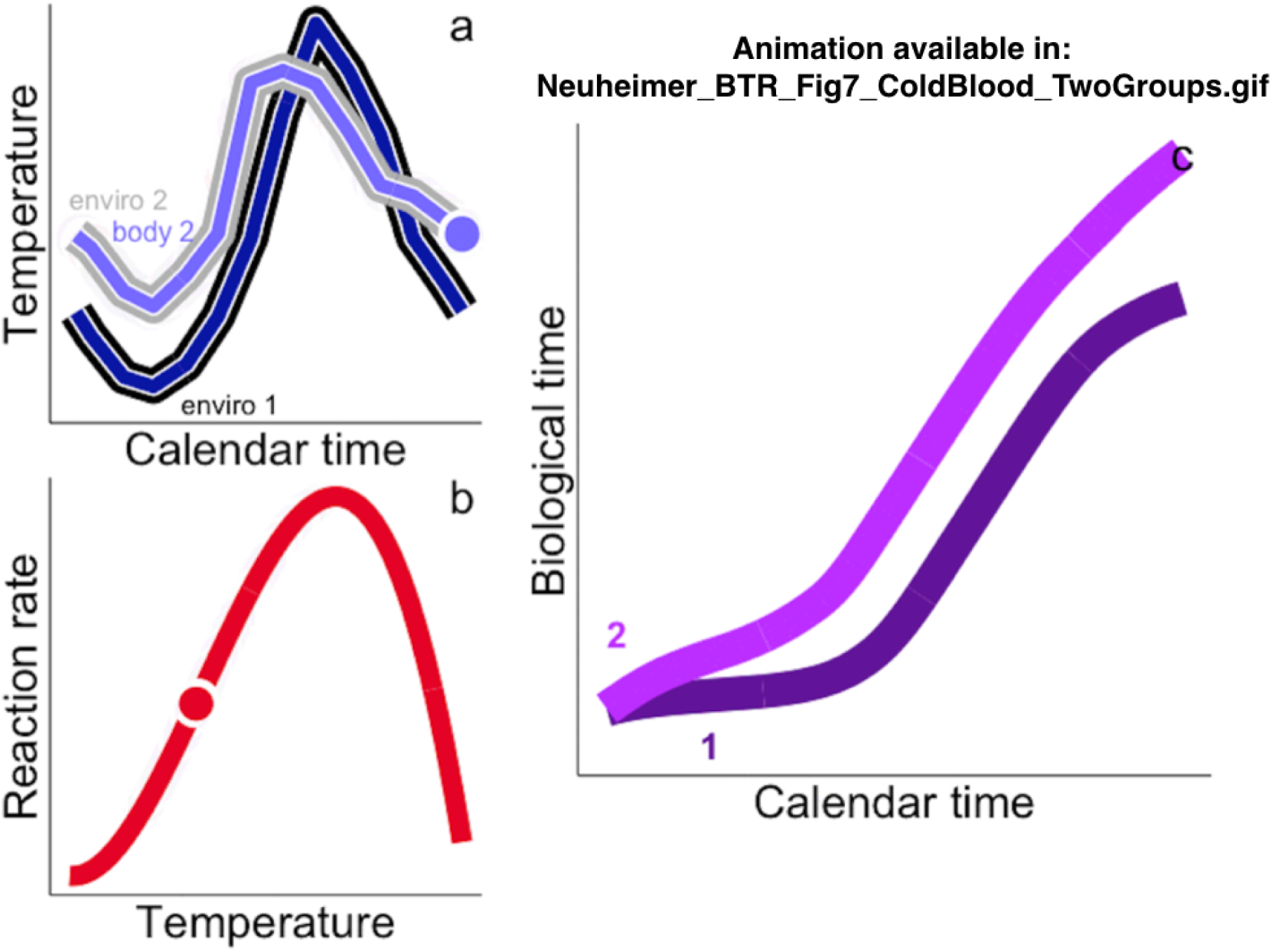
How biological time compares across groups for cold-blooded organisms. a) Environmental temperatures changing over space or years (grey vs. black) results in changes to the organism’s body temperature (light vs. dark blue). b) Body temperature variations changes the scaling from calendar time to biological time via the TPC. c) As a result, biological time accumulates differently for cold-blooded organisms across space or time (light vs. dark purple).

**Fig. 8:**
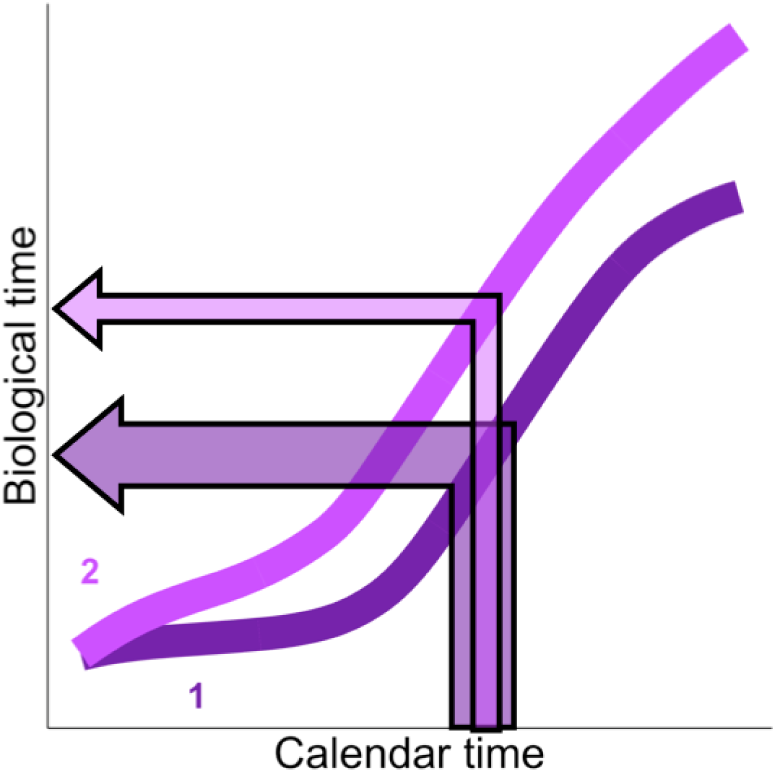
Observations of cold-blooded organisms across groups (e.g. cohorts or locations, light purple vs. dark purple) are only directly comparable once converted to relevant biological time-scales (arrows).

The difference between how biological time emerges for warm-vs. cold-blooded organisms sets up the risk of our “warm-blooded bias” making patterns in biological observations difficult to see and/or misinterpretted when viewed only through a calendar time frame. Luckily, the BTR both identifies the problem and allows us to overcome our bias by translating the time-scales of observations between the two temporal frames of reference. Doing so, both clarifies the influence of temperature on observed biological variability, and allows us to disentangle temperature-dependent variability from that due to other factors (e.g. food, size-structure, fishing, trade-offs across processes; e.g. [13]).

The idea of estimating a thermally dependent biological time-scale to explain variability originated at least 285 years ago when the French naturalist Réaumur (1735; [14]) put forward the idea of “degree-days” (DD) as a simple sum of the available heat energy (temperature) during growth and development:

> *“The same grains are harvested in very different climates; it would be interesting to compare the sums of heat degrees* [biological time] *over the months during which wheat does most of its growing and reaches complete maturity in hot countries, like Spain or Africa … in temperate countries like France … and in the colder countries of the North.”* Original text in Old French: [15]; Quoted in [16].

Since Réaumur, biological time as degree-days, DD, (or “growing degree-days”, GDD) has been used in agriculture to estimate growing season, plan crop harvest timing, etc. (reviewed in [16]). Similarly, entomologists (beginning at least 70 years ago) began exploiting biological time to explain growth and development in insects (e.g. [17]). More recently (and formally since [18]), the use has been expanded to aquatic animals with size-at-age and development timing explained on biological time for fish and invertebrates (e.g. [18], [19], [20] [21], [22], [23] [24], [25]). For example, we were able to use DD as a biological (thermal) time-scale that explains 93% of the variability in juvenile cod (*Gadus morhua*) size-at-age across the species’ range [18], as well as disentangle adaptive variability in reproductive timing across years and locations [21], [25].

Metrics used to capture the relevant biological time-scale have evolved over the years along with more detailed characterizations of the TPC. Réaumur (1735; [14]) describes degree-days as a simple sum of temperatures above a threshold, a metric that assumes we are in the mid-range of temperatures where the TPC is near-linear (Fig. 1). In contrast, more recent studies show the power of including the non-linear nature of the TPC to accurately measure biological time with methods using nonlinear averaging (e.g. [26], [27]) and fractional development techniques (e.g. MCF, [28], [29], [25]), to represent the TPC scaling of calendar time. For example, Bernhardt et al. (2018; [26]) predict growth rates of a green alga (*Tetraselmis tetrahele*) with high accuracy using a non-linear averaging method that respects the non-linear nature of the TPC. The high explained variability and predictive power demonstrates the significance and usefulness of the BTR - patterns become clear in the noise as we develop biological time metrics that incorporate the mechanistic foundation underlying the relationship between biological and calendar time (scaling via the TPC).

#### Predicting the world around us

The previous section showed how an understanding of the BTR allows us to assign a relevant biological time-scale to observations made in calendar time. In so doing, we clarify observed variability in timing and are able to assign the variability to temperature dependent vs. independent forcing factors. When we move from calendar time to biological time in this way, we identify relevant biological time-scales that can be thought of as “thermal constants” [18] representing the amount of biological time (time scaled by temperature via the TPC) that is required to reach a given biological threshold (e.g. moving from stages I to II in Fig. 9). Combined with knowledge of how an organism’s biological time accumulates, thermal constants can be used to predict the calendar timing of biological phenomena: a biological event is expected to be observed in calendar time when an organism’s relevant biological time intersects the thermal constant (Fig. 9). In this way, we can move from a biological time scale back to calendar time to estimate when a biological phenomenon (emergence time, time to stage) is likely to be observed, and predict how that timing will interact with timing of other processes (e.g. growth vs. development) or organisms (e.g. across trophic levels), including ourselves (see also Mapping out the BTR research landscape below).

**Fig. 9:**
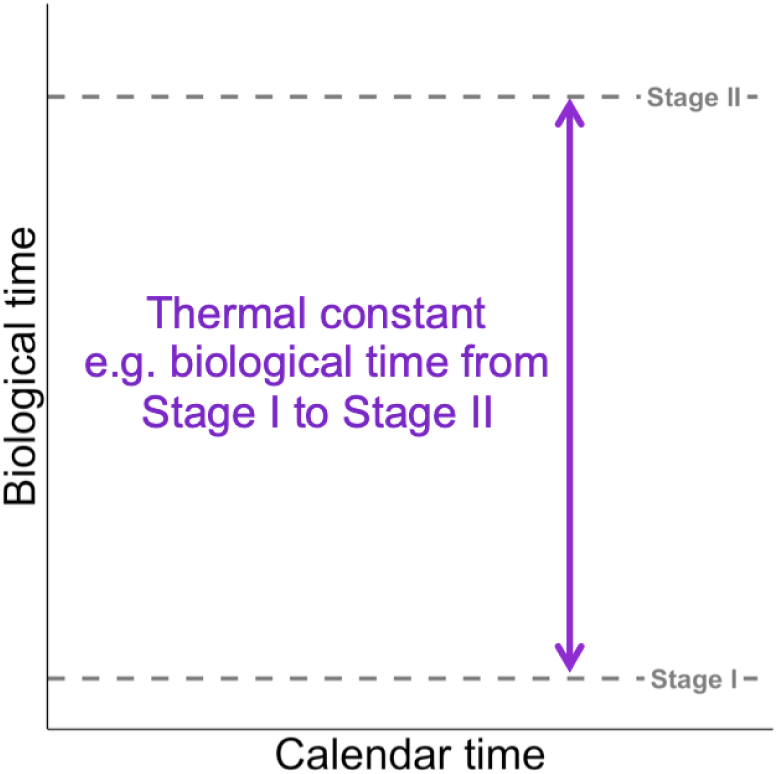
Thermal constants describe the biological time (time scaled by temperature as described in the BTR) to accomplish a biological threshold, e.g. moving from a theoretical Stage I to Stage II.

Again, let’s explore how we can use the idea of thermal constants to predict the timing of events in calendar time for warm-vs. cold-blooded organisms: For warm-blooded organisms, biological time accumulates at a constant rate that is consistent across space or time (e.g. groups 1 and 2 in Fig. 10). Assuming food, etc. are constant across groups (years or locations; see Explaining the world around us), biological time accumulates similarly across the groups of warm-blooded animals and the trajectory of biological time intersects the thermal constant such that the timing of events are seen as consistent in both calendar and biological time (Fig. 10). Thus, all else being equal (including body temperatures, the TPC and temperature independent factors such as food), predicted biological timing will appear unchanged across years or locations for warm-blooded organisms.

**Fig. 10:**
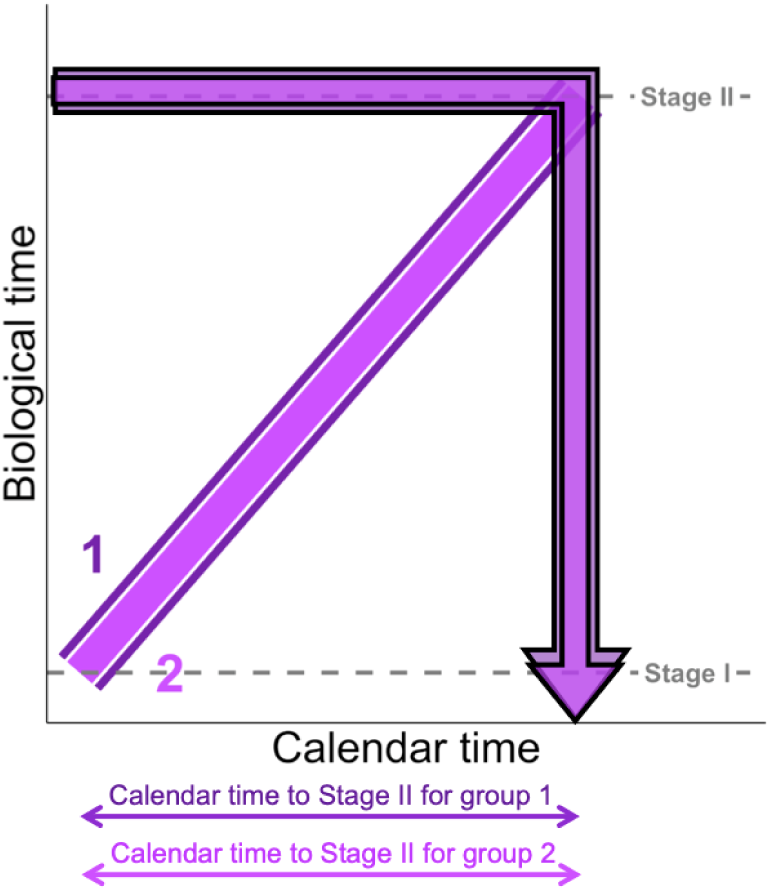
Using a thermal constant to move from a biological time-scale to a calendar time-scale and estimate time to Stage II for different groups (e.g. cohorts or locations) of a warm-blooded organism.

Now, let’s explore how thermal constants will affect observations of cold-blooded timing across groups (years or locations). Again, we define the thermal constant as the biological time needed to meet a biological threshold, e.g. to advance from Stage I to Stage II (Fig. 11). When environmental temperatures differ among groups of cold-blooded animals, biological time accumulates differently relative to calendar time. These different time trajectories result in intersections with the thermal constant that vary in calendar time across the groups (Fig. 11), and thus, the predicted calendar time occurence of the phenomenon (i.e. advancing to Stage II) varies across the groups (Fig. 11). Again, other factors may also influence predicted timing (e.g. food consumption, stage-structure, adaptation of TPC) - but accurate predictions as well as the influence of these other factors can only be estimated when we begin with predictions that account for the relative nature of biological time through the BTR.

**Fig. 11:**
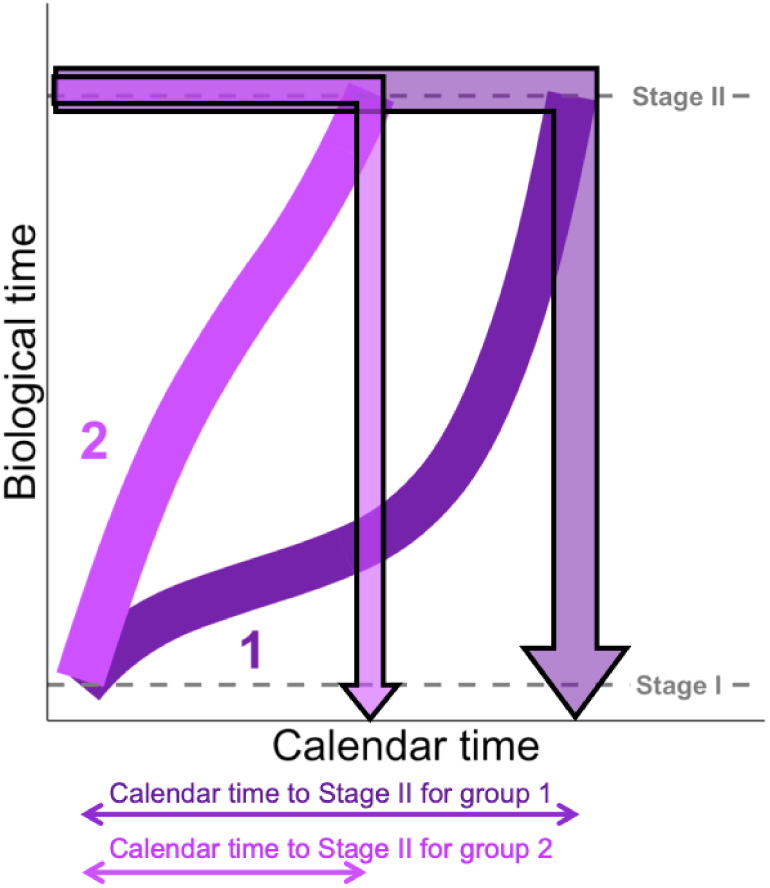
Using a thermal constant to move from a biological time-scale to a calendar time-scale and estimate time to Stage II for different groups (e.g. cohorts or locations) of a cold-blooded organism.

Identifying relevant thermal constants in this way allows us to both hindcast the timing of unobserved phenomena (e.g. larval stages), and forecast timing predicted under future temperature conditions. Applications of the BTR in this way are found in previous work estimating the timing of unobserved stages of larval fish and their prey across populations (e.g. [25]), and predicting future reproductive timing dynamics under climate change scenarios (e.g. [21]).

The idea of using biological clocks to predict calendar timing of biological events is also at the heart of a series of Hawaiian sayings, or Ōlelo no eau, originating over centuries (compiled in [30]) that describe timings of seasonal biological events (e.g. flowering, arrival, etc.) synchronized across organisms (e.g. Box 1). These sayings often link timing between terrestrial and aquatic players and were used to e.g. predict harvest time, the arrival of seasonal dangers, etc. In each saying, the timing of one organism (e.g. plant flowering) is used to predict the timing of another (e.g. arrival of a fish), with the conincidence of the two phenomena maintained from year to year despite possible shifts in the organisms’ respective environments. The basis of the coincidence can be explained by an underlying similarity in the pace of the species’ respective biological time-scales resulting from a combination of their respective body temperatures scaled through their respective TPCs (i.e. *f*(*T*(*t*)) in eqn (3)). While calendar time is unable to capture the year to year changes in the pace of these “biological clocks”, the sayings allow us to appropriate the biological clock of one phenomenon (usually a relatively easy to observe terrestrial organism, e.g. flowering plant) to predict the timing of another (usually a less easy-to-observe aquatic organism, e.g. arrival of a fish).

##### Box 1: Examples of Hawaiian sayings (or ‘Ōlelo no ‘ eau) describing links in biological timing across species

**Pala ka hala, momona ka wana.**

*When the hala is ripe, the urchin is fat*

When the fruit of the hala tree (*Pandanus tectorius*) is ripe, the sea Urchin is full of gonads.

**Pua ke kō, ku ka he’ e.**

*When the sugar cane tassels, it is octopus season.*

When the sugar cane is, mature, the octopus are in the coastal waters.

**Pua ka wiliwili, nanahu ka manō.**

*When the wiliwili tree blooms, the sharks bite.*

The timing of the wiliwili tree *(Erythrina sandwicensis)* blooming, Coincides with when tiger shark *(Galeocerdo cuvier)* are birthing their young, and the mothers are more aggressive.

#### Manipulating the world around us

An understanding of the BTR also gives us the opportunity to shape the timing of biological events in calendar time for convenience, economy, etc. A familiar example of this comes from popular food storage methods where lowering temperature allows us to manipulate (slow) the pace of biological time to delay food spoiling (Fig. 12). By cooling the temperature in the food’s immediate environment (e.g. putting milk in the fridge; Fig. 12), we change the pace at which biological time of e.g. microbial growth is accumulating with respect to calendar time, and lengthen the (calendar) time needed to reach the thermal constant of food spoiling (Fig. 12). Similar manipulations of biological time are already in use and widely familiar both in the home (e.g. raising the temperature to shorten the calendar time needed for dough to rise, or placing cut flowers in a cool room to lengthen their calendar time lifespan) as well as commercially (e.g. to manipulate crops in agriculture and aquaculture).

**Fig. 12:**
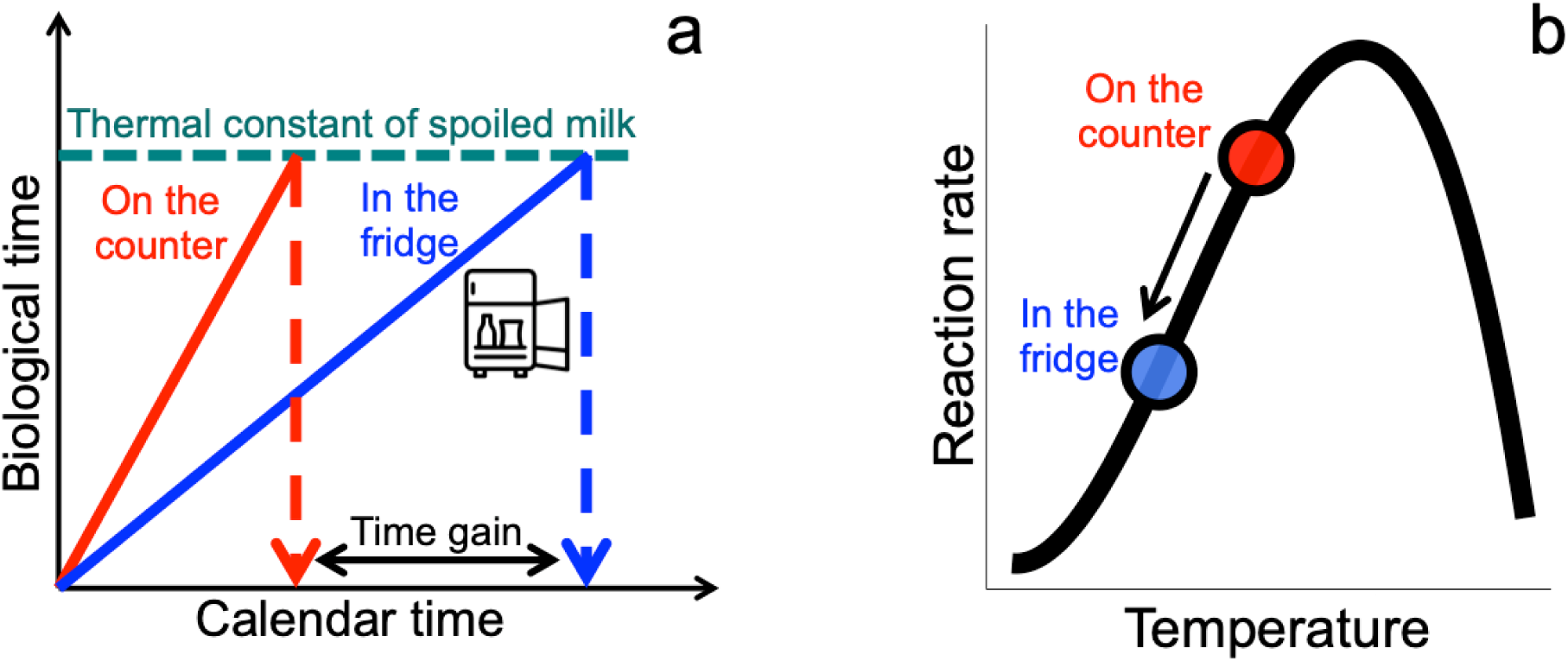
An example of how the BTR helps us manipulate biological phenomena: The biological time to milk spoiling can be represented as a thermal constant (a). When the biological age of the milk intersects this thermal constant, the milk is spoiled. We can manipulate the calendar time when this intersection occurs: by putting the milk in the fridge, we lower the “body” temperature of the microbes involved in the spoiling (b). This lower temperature means slower reaction rates (b) and the scaling of biological time relative to calendar time is changed. Biological age for milk in the fridge accumulates more slowly than milk left on the counter, and the intersection with the thermal constant of spoiled milk happens later in calendar time (a). Icon by Vectors Market from the Noun Project.

To summarize this section, there is great utility to be found in the BTR and its description of the relationship between calendar and biological temporal frames of reference. Understanding how biological time progresses relative to calendar time allows us to clarify and explain observed variability and identify typical biological time-scales associated with biological events (thermal constants). This latter tool can be used to both make predictions of unobserved timings, in the past or future [25], as well as to manipulate the timing of the biological world around us. These opportunities present a landscape of research opportunities in biology and associated fields - this is the subject of the following section.

### Mapping out the BTR research landscape

The BTR presents a natural framework to shape useful biological questions and future work. These questions can be structured around the BTR equation (eqn (3) above) - covering topics related to body temperatures, thermal performance curves, and the resulting biological time-scales themselves.

#### Body temperature

An understanding of biological time-scales requires accurate descriptions of body temperature of the target process or organism. Important questions include:

- How correlated are relevant body temperatures to the environment?
- How do body temperatures change in space and time?
- Why and how do cold-blooded animals manipulate their body temperature? (See How biological time scales with temperature)

#### Thermal Performance Curve (TPCs)

How biological time accumulates relative to calendar time requires characterization of how time scales with different temperatures described by the thermal performance curve(s) or TPCs. Relevant questions include:

- How do TPCs change across individuals, time or space? Here lies questions of (intra-species) adaptation.
- When can TPCs be generalized across groups?
- How are TPCs affected by other factors such as size or food?
- How do TPCs interact across processes (growth, development) and what are the resulting effects in calendar time? Here lies the basis of the Temperature-Size Rule ([31]).

#### Exploring life on biological time

As described in Using the biological theory of relativity above, combining information on body temperature and TPC allows us to estimate biological time-scales via the BTR. We are then positioned to tackle a number of fundamental questions including:

- Exploring biological variability on biological time:
  – How do trends in size, maturity, lifespan, etc. vary on relevant biological time-scales? e.g. [18], [19], [20], [22].
  – What variability remains when observations are positioned in biological time, and how can it be explained by other factors, e.g. light, food, fishing, etc.? e.g. [32], [13].
  – How is the timing of biological phenomena likely to change over space and time? e.g. [21], [33], [34].
- Species interactions:
  – How do timing controls (thermal or otherwise) vary across species including trophic levels? e.g. [25].
  – How do predators predict prey timing when the pace of biological time differs across species? e.g. herbivores feeding on plants where plant timing is limited by seasonal light availability, warm-blooded mammals predicting the timing of seasonal cold-blooded prey (e.g. whales and krill).
  – How can humans translate our behaviour onto relevant biological time-scales to better strategize our interactions with other biological players? e.g. basing levels of exploitation on relevant thermal years (rather than constant calendar years), or converting survey sample dates to an equivalent biological time-stamp to avoid errors (e.g. aliasing) when comparing observations across space or time [35].

## Conclusion

In summary, I present a theory of relativity for biology that allows us to move across temporal frames of reference to gain a relevant measure of time for biological processes and players. The BTR allows us to describe how biological time emerges relative to calendar time, and observe biological variation on an organism’s own time-scale. In a manner similar to Einstien’s STR, this change of reference frame allows us to explain observed changes in space and time, and make predictions of biological variation into the future. Moreover, we can use the BTR to engineer/tailor our interactions with the biological world to increase the economic and/or conservation success of our actions. In these ways, the BTR offers both a mechanistic explanation and a translation tool to move between temporal frames of reference to better understand the biological world.

## Supporting information

Figure 3 as animation

Figure 4 as animation

Figure 5 as animation

Figure 7 as animation

Manuscript at html with embedded animations

## Acknowledgements

This work was completed during an AIAS-COFUND Fellowship at the Aarhus Institute of Advanced Studies, which receives funding from the Aarhus University Research Foundation (Aarhus Universitets Forskningsfond) and the European Union’s Seventh Framework Programme, Marie Curie Actions (grant agreement 609033). ABN is grateful to Rosie Alegado, Kyle Edwards, Kumu Hula Māliā Helelā, Karen Marie Hilligsøe, and Luís P. Viegas for conversations that helped clarify the content of this manuscript. Manuscript and animations made with [36], [37] and [38] in [39]. This work was completed in part at the North Market Cafe in Almonte, Ontario, Canada, and ABN is grateful for their hospitality.

