## Supplementary figures and images for "The pace of life: Time, temperature, and a biological theory of relativity"

### Figure 3 as animation

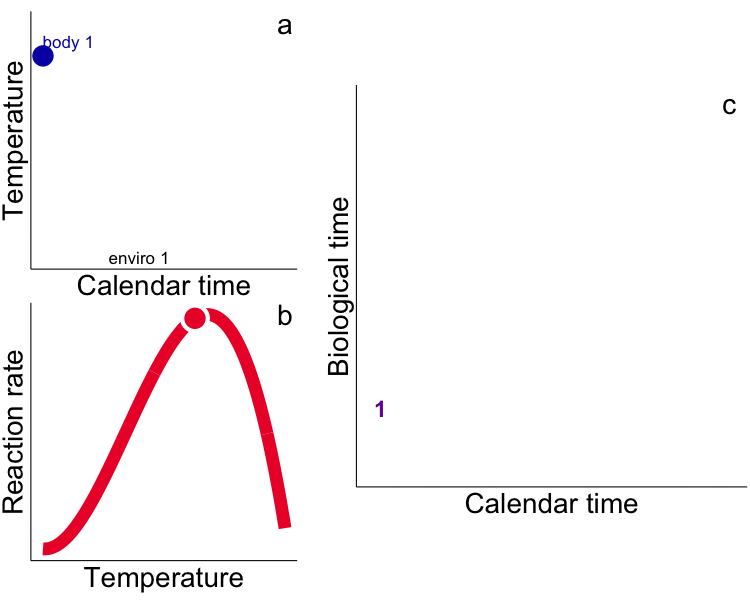

### Figure 4 as animation

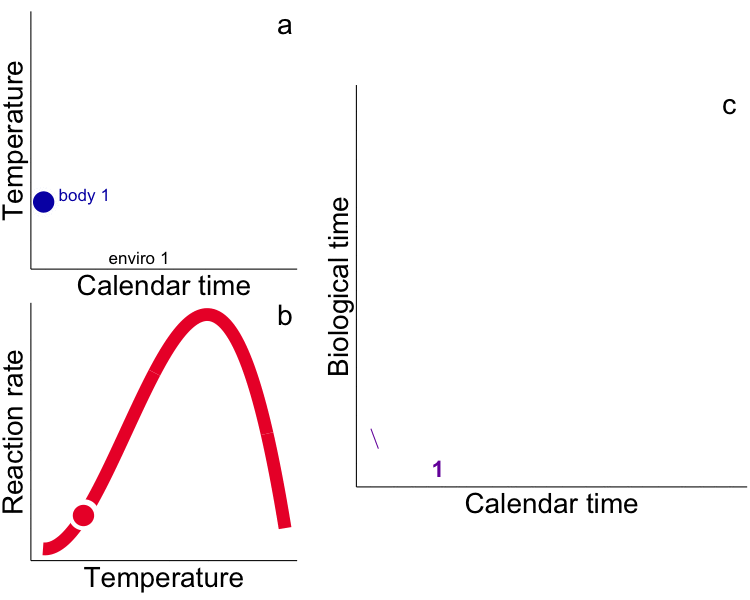

### Figure 5 as animation

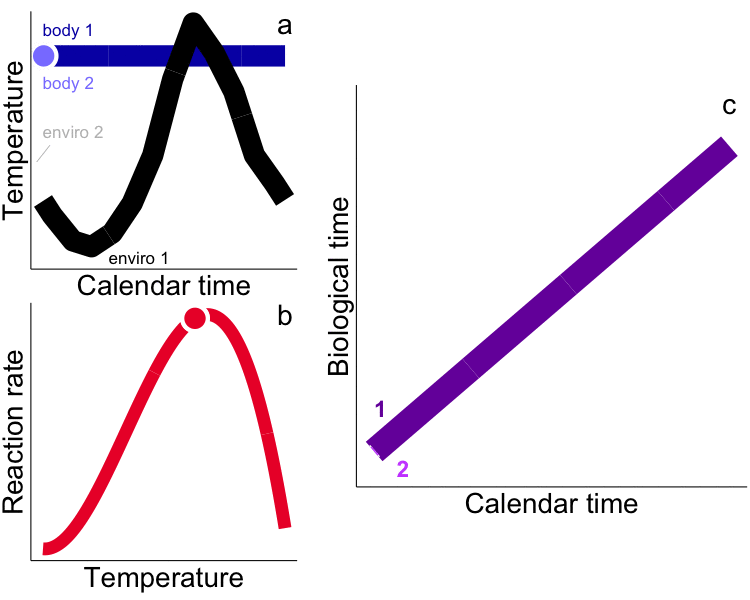

### Figure 7 as animation

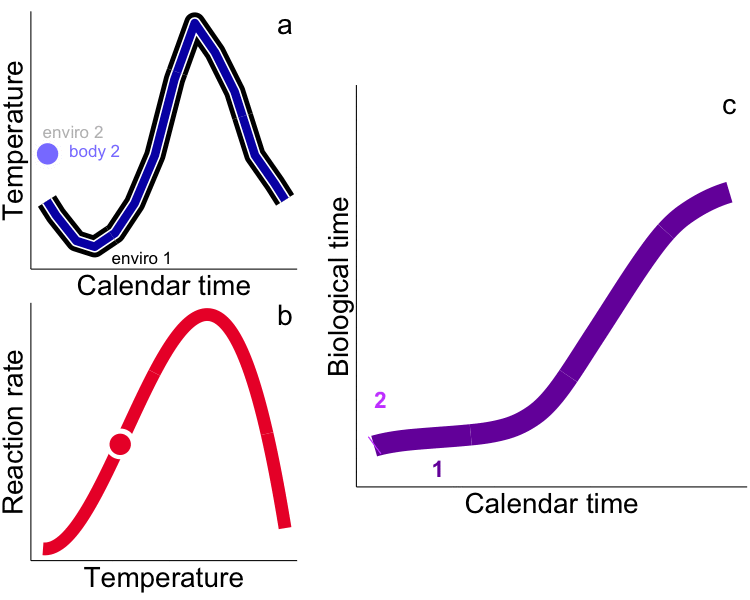
