## Supplementary material for "The pace of life: Time, temperature, and a biological theory of relativity": Manuscript at html with embedded animations: Neuheimer_BioTheoryOfRelativity_BioRxiv_Manuscript_HTML_Correction01.html

Neuheimer\_BioTheoryOfRelativity\_BioRxiv\_Manuscript\_HTML\_Correction01.utf8

Anna B. Neuheimer*^1,2^*

*^1^Aarhus Institute of Advanced Studies (AIAS), Aarhus University, DK-8000 Aarhus C, Denmark*  
*^2^Department of Oceanography, University of Hawaiʻi at Mānoa, 1000 Pope Road, Honolulu, HI 96822, USA*

Fig. 2: Parallels between the Special Theory of Relativity and the Biological Theory of Relativity. See text for description.

The STR describes how time scales with velocity:

\[\begin{equation}
\Delta t\_{mov} = f(v)·\Delta t\_{rest}
\tag{1}
\end{equation}\]

where \(\Delta t\_{mov}\) is the ellapsed time in the moving frame determined by the ellapsed time at rest \(\left(\Delta t\_{rest}\right)\) and a velocity \(\left(v\right)\) scaling \(\left(f(v)\right)\), i.e. the Lorentz factor \(\left(\gamma = \frac{1}{\sqrt[\leftroot{-2}\uproot{5}]{1-\frac{v^2}{c^2}}}\right)\) where \(c\) is the speed of light in a vacuum.

Similarly, the BTR describes how biological time scales with temperature as:

\[\begin{equation}
\Delta t\_{bio} = f(T)·\Delta t\_{cal}
\tag{2}
\end{equation}\]

where \(\Delta t\_{bio}\) is biologically relevant elapsed time (i.e. in the biological frame of reference) determined by the elapsed “calendar” time \(\left(\Delta t\_{cal}\right)\) scaled with the relevant body temperature \(\left(T\right)\) via the TPC \(\left(f(T)\right)\). Both body temperature \(\left(T\right)\) and the TPC \(\left(f(T)\right)\) may change in time & space (including across individuals). I will cover what happens when the TPC scaling changes in Mapping out the BTR research landscape below, but here let’s expand eqn (2) to allow for changes in body temperature over time \(\left(T(t)\right)\):
